# Tracking the embryonic stem cell transition from ground state pluripotency

**DOI:** 10.1101/092510

**Authors:** Tüzer Kalkan, Nelly Olova, Mila Roode, Carla Mulas, Heather J. Lee, Isabelle Nett, Hendrik Marks, Rachael Walker, Hendrik G. Stunnenberg, Kathryn S. Lilley, Jennifer Nichols, Wolf Reik, Paul Bertone, Austin Smith

**Affiliations:** Wellcome Trust-Medical Research Council Cambridge Stem Cell Institute, Cambridge CB2 1QR, UK; Babraham Institute, Cambridge CB22 3AT, UK; Radboud University, Faculty of Science, Department of Molecular Biology, Radboud Institute for Molecular Life Sciences (RIMLS), 6500HB, Nijmegen, The Netherlands; Department of Physiology, Development and Neuroscience, University of Cambridge, Cambridge CB2 4BG; Department of Biochemistry, University of Cambridge, Cambridge CB2 1GA; The Cambridge Centre for Proteomics, Cambridge System Biology Centre, University of Cambridge, Cambridge, CB2 1QR, UK; Wellcome Trust Sanger Institute, Hinxton CB10 1SA, UK; Centre for Trophoblast Research, University of Cambridge, Cambridge CB2 3EG, UK

## Abstract

Mouse embryonic stem (ES) cells are locked into self-renewal by shielding from inductive cues. Release from this ground state in minimal conditions offers a system for delineating developmental progression from naive pluripotency. Here we examined the initial transition of ES cells. The population behaves asynchronously. We therefore exploited a short-half-life *Rex1::GFP* reporter to isolate cells either side of exit from naive status. Extinction of ES cell identity in single cells is acute. It occurs only after near-complete elimination of naïve pluripotency factors, but precedes appearance of lineage specification markers. Cells newly departed from the ES cell state exhibit global transcriptome features consistent with features of early post-implantation epiblast and distinct from primed epiblast. They also exhibit a genome-wide increase in DNA methylation, intermediate between early and late epiblast. These findings are consistent with the proposition that naïve cells transition to a discrete formative phase of pluripotency preparatory to lineage priming.

**Highlights:**

- The Rex1 destabilized GFP reporter demarcates naive pluripotency.
- Exit from the naive state is asynchronous in the population.
- Transition is relatively acute in individual cells and precedes lineage priming.
- Transcriptome and DNA methylome reflect events in the pre-gastrulation embryo.

## Introduction

Epiblast cells, founders of all somatic cells and the germ line, are formed in the inner cell mass (ICM) in the final day of pre-implantation development in mice (Boroviak and Nichols, 2014; Cockburn and Rossant, 2010). This emergent condition of “naive pluripotency” (Nichols and Smith, 2009) is characterized by a unique suite of transcription factors, a hypomethylated genome, and the ability to give rise directly and clonally to embryonic stem (ES) cells (Boroviak et al., 2014; Brook and Gardner, 1997; Lee et al., 2014; Smith et al., 2012). Upon implantation, ES cell forming capacity is abruptly lost, epithelial organisation becomes manifest, global gene expression is reconfigured, and DNA methylation increases within 24 hours, indicative of a profound cellular transition (Auclair et al., 2014; Bedzhov and Zernicka-Goetz, 2014; Boroviak et al., 2014; Boroviak et al., 2015). Nonetheless, post-implantation epiblast cells retain pluripotency (Osorno et al., 2012; Solter et al., 1970). However, leading up to gastrulation egg cylinder epiblast cells are subject to inductive cues and become fated, though not yet lineage-committed (Osorno et al., 2012; Solter et al., 1970; Tam and Zhou, 1996).This late phase of pluripotency during gastrulation is termed primed, reflecting the incipient expression of lineage specification factors (Hackett and Surani, 2014; Nichols and Smith, 2009).

Mouse ES cells cultured in serum-free media supplemented with chemical inhibitors of MEK1/2 and GSK3α/β, known as ‘2i’ (Ying et al., 2008), and optionally the cytokine LIF are in a uniform condition of self-renewal termed the “ground state”. Ground state ES cells show transcriptional and epigenetic similarity to naive pre-implantation epiblast (Boroviak et al., 2014; Ficz et al., 2013; Habibi et al., 2013; Leitch et al., 2013; Nichols and Smith, 2012). Upon withdrawal of 2i, ES cells embark on a path to lineage commitment (Dunn et al., 2014; Marks et al., 2012; Ying et al., 2008). Recent studies have begun to explore the dissolution of naive pluripotency and the route towards multi-lineage differentiation *in vitro* (Acampora et al., 2013; Betschinger et al., 2013; Buecker et al., 2014; Davies et al., 2013; Kurimoto et al., 2015; Leeb et al., 2014; Liu et al., 2015; Respuela et al., 2016; Thomson et al., 2011; Yang et al., 2014). However, in contrast to the homogeneity of ES cells in 2i/L, differentiating cultures become heterogeneous (Buecker et al., 2014; Hayashi et al., 2011; Kalkan and Smith, 2014; Marks et al., 2012). A means to identify and select cells as they transition from naive pluripotency would facilitate experimental resolution.

We previously generated ES cells carrying a *Rex1::GFPd2* (RGd2) reporter in which the coding sequence of one allele of Rex1 (gene name *Zfp42)* is replaced with a GFPd2-IRES-*bsd* cassette that produces a destabilized version of GFP protein with a 2-hour-half-life (GFPd2) (Wray et al., 2011). Here we exploit this reporter to monitor exit from naive pluripotency for mouse ES cells guided by autocrine cues in defined adherent culture. We test the utility of the reporter as a faithful marker of naive pluripotency and survey transcriptional and DNA methylome changes during the initial transition towards differentiation competence.

## Results

### Pluripotency is unaffected by the RGd2 reporter knock-in

The Rex1 coding sequence is entirely deleted in the *RGd2* allele. RGd2 ES cells (Wray et al., 2011) were transmitted through the mouse germline and heterozygous animals were backcrossed twice to strain 129. Following heterozygous intercrosses, homozygous mice were healthy and fertile, although slightly under-represented (Table S1). These results confirm previous reports that Rex1 is dispensable for development (Kim et al., 2011; Masui et al., 2008; Rezende et al., 2011). We could derive wildtype, heterozygous and homozygous ES cells, both male and female, from intercross embryos (Table S2), demonstrating that Rex1 is not important for ES cell propagation. RGd2 expression should therefore constitute a neutral reporter.

We evaluated reporter expression in the embryo. Dual immunofluorescence staining for GFP and Nanog revealed that the RGd2 reporter is expressed uniformly expressed throughout the naive epiblast at E4.5, whereas Nanog expression is heteregenous (Fig 1). Both were excluded from the primitive endoderm (PrE) and trophoblast. Shortly after implantation at E5, neither Nanog nor GFP are detected in the epiblast, whereas GFP appears in the extra-embryonic ectoderm at this stage (Fig 1). These results are consistent with in situ RNA hybridisation (Pelton et al., 2002) and quantitation of Rex1 mRNA expression (Boroviak et al., 2015) during epiblast progression. We conclude that the *RGd2* allele faithfully reports endogenous *Rex1* transcription and accordingly that GFP expression coincides with naive pluripotency *in vivo* (Boroviak et al., 2014).

**Fig 1.**
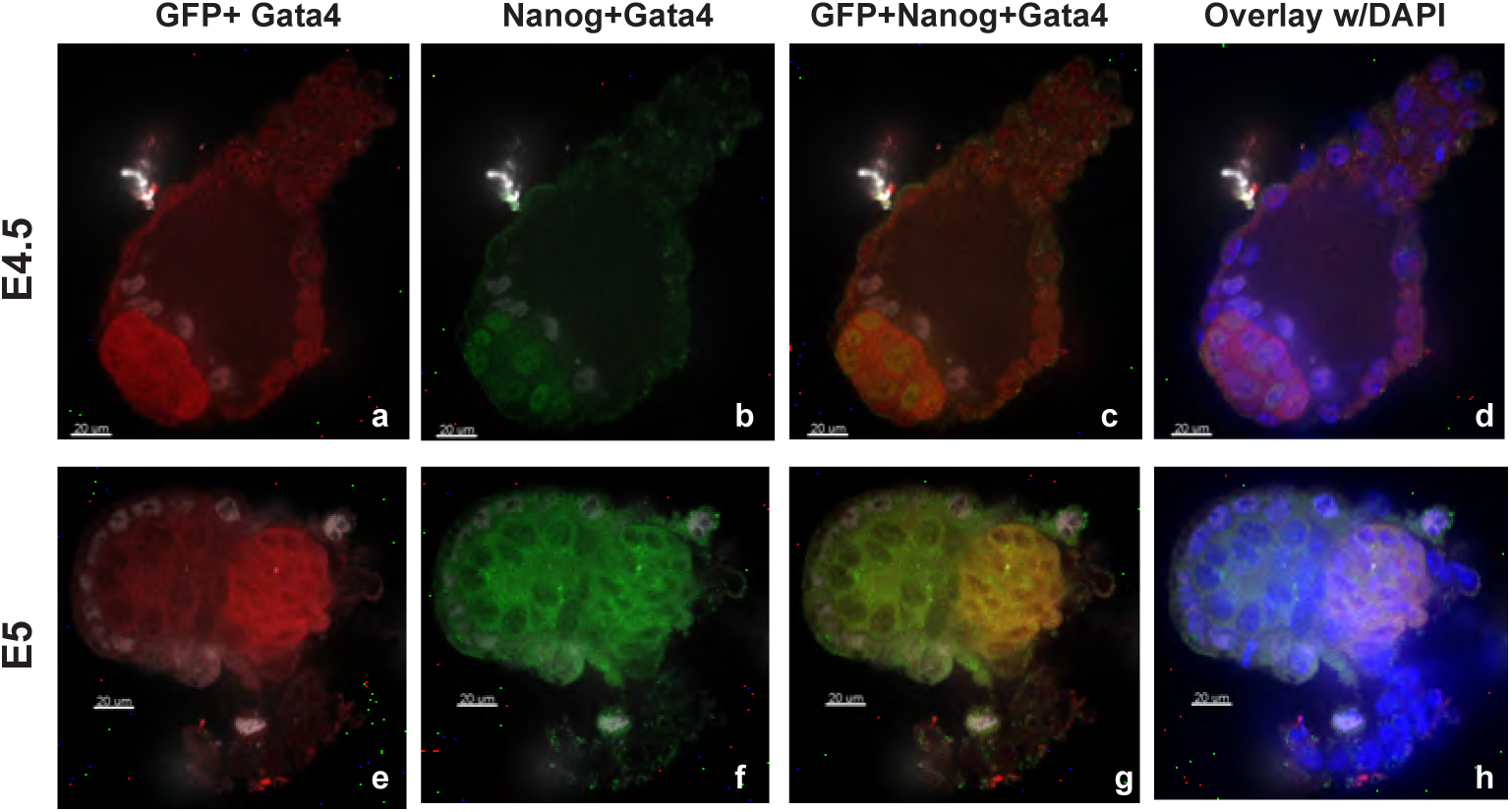
Expression of RGd2 reporter in the embryo. **(A)** IF staining for GFP (Rex1GFPd2) (red), Nanog (green), Gata4 (gray), DAPI (blue) at E4.5 (a-d) and E5 (e-h).Scale bar is 20μm.

### Release of ES cells from 2i triggers progression towards multi-lineage specification

We monitored the early phase of ES cell transition *in vitro* after withdrawal from 2i in serum-free N2B27 medium on gelatin-coated plastic (Fig 2A). We started from ES cells in 2i alone because LIF delays the onset of differentiation (Dunn et al., 2014). Plating ES cells directly in N2B27 at low density (<10000 cells cm-^2^) results primarily in neural specification (Ying et al., 2003). However, when ES cells were plated at an intermediate density (15000 cells cm-^2^) and maintained in 2i for 24 h prior to 2i withdrawal, numerous Brachyury (*T*) positive cells also appeared, indicative of non-neural fates (Fig 2B). These conditions were used throughout this study.

**Fig 2.**
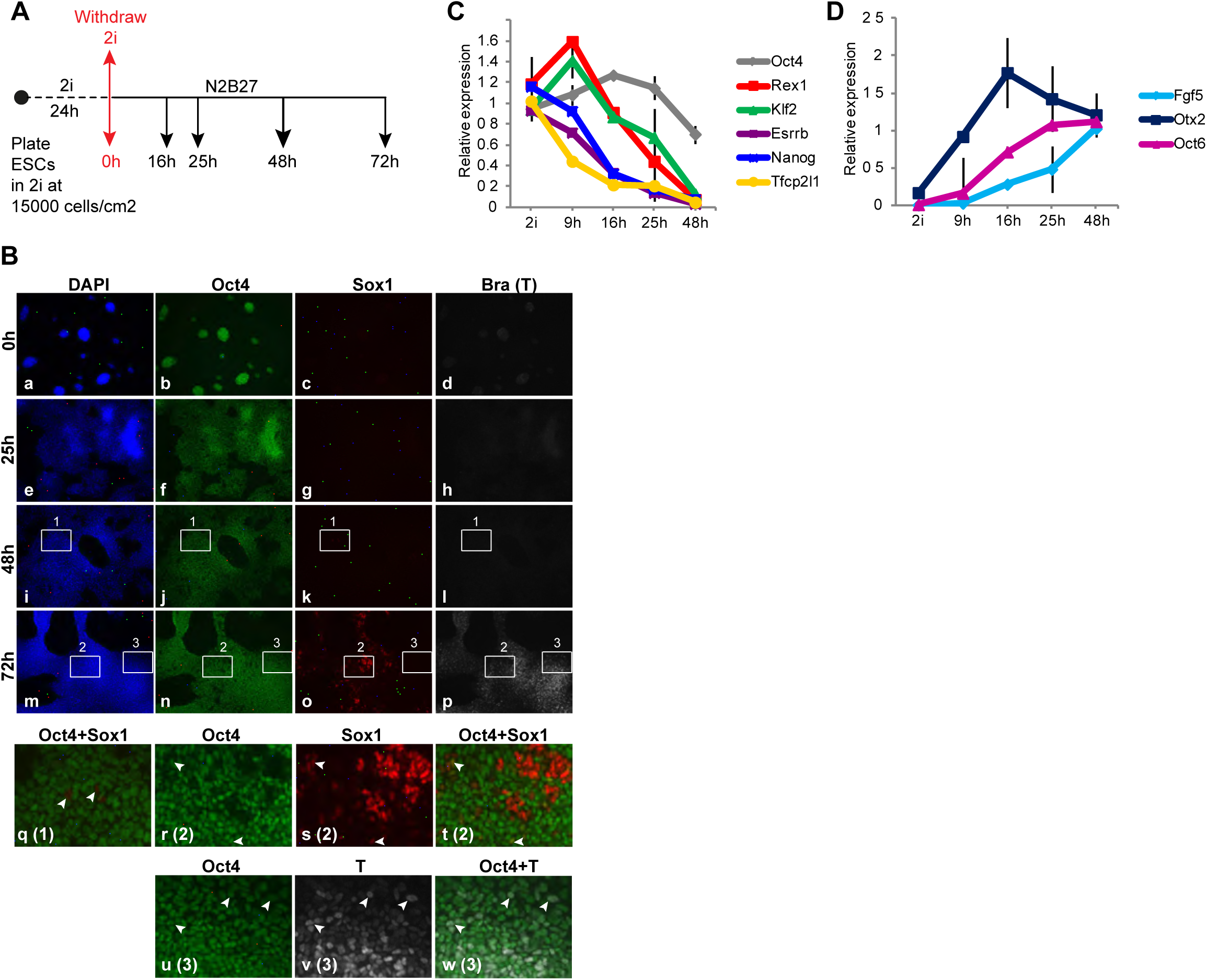
Multilineage differentiation of ES cells upon release from 2i. **(A)** Experimental protocol for monolayer differentiation of naive ES cells in N2B27 by withdrawal of 2i. **(B)** IF staining for Oct4 (green), Sox1 (red) and Bra (*T*) (gray) at the indicated time points post-2i withdrawal. Panels **q** through **w** show enlarged insets from 48 h and 72 h, with respective inset number in parantheses. q-RT-PCR for selected **(C)** pluripotency and **(D)** early post-implantation epiblast markers. Expression levels are shown as fold change relative to naive ES cells in **C** and 48h samples in **D**. GAPDH was used for normalization of cDNA levels.

Protein expression of Oct4 was relatively uniform for 48 h after release from 2i (Fig 2B). Rare cells expressing low levels of Sox1 were first detected at 48h (Fig 2B; i-l, arrowheads in q). By 72 h clusters of bright Sox1-positive cells that lacked Oct4 emerged. Occasional Sox1/Oct4-double-positive cells were found outside these clusters (Fig 2B; arrowheads in r-t). T-expressing cells were first detected at 72 h, mostly in dense clumps that were mutually exclusive with Sox1-positive clusters (Fig 2B, m-p). T-positive cells at this stage were also positive for Oct4 (Fig 2B; arrowheads in u-w), consistent with transient co-expression in early primitive streak and during directed *in vitro* differentiation (Hoffman et al., 2013; Thomson et al., 2011).

Oct4 mRNA was mildly downregulated only at 48 h (Fig 2C). In contrast, transcripts for naive transcription factors (TFs) (Dunn et al., 2014; Martello and Smith, 2014) declined within the first 25 h. Downregulation of Nanog, Esrrb and Tfcp2l1 initiated before Rex1 and Klf2. Concurrently, transcripts for post-implantation epiblast markers Fgf5, Otx2, Oct6 (*Pou3f1*), were upregulated (Fig 2D). Transcripts for naive TFs were almost eliminated at 48h (Fig 2C). Similar results were observed with multiple ES cell lines (Fig S1A, B). These results indicate that upon release from self-renewal, in defined conditions driven by autocrine signals ES cells progress from the naive state to multi-lineage specification in an orderly sequence. First naive TFs are extinguished and markers diagnostic of post-implantation epiblast are induced. Subsequently lineage specific markers emerge and Oct4 is downregulated.

### Pluripotency factors display individual downregulation dynamics upon 2i withdrawal

To follow the kinetics of transition following release from 2i, we monitored the RGd2 reporter by flow cytometry (Fig 3A). GFP was expressed unimodally with a narrow distribution in 2i. This tight peak persisted throughout the first 16h after withdrawal, although mean fluorescence rose slightly. Rex1 mRNA did not increase (Fig 2C), therefore this shift may result from an increase in translation or GFP half-life. By 25 h, GFP intensity became heterogeneous with many cells shifted to lower expression. This suggests that Rex1 may be downregulated with different kinetics in individual cells. By 48 h the majority of cells had extinguished GFP.

**Fig 3.**
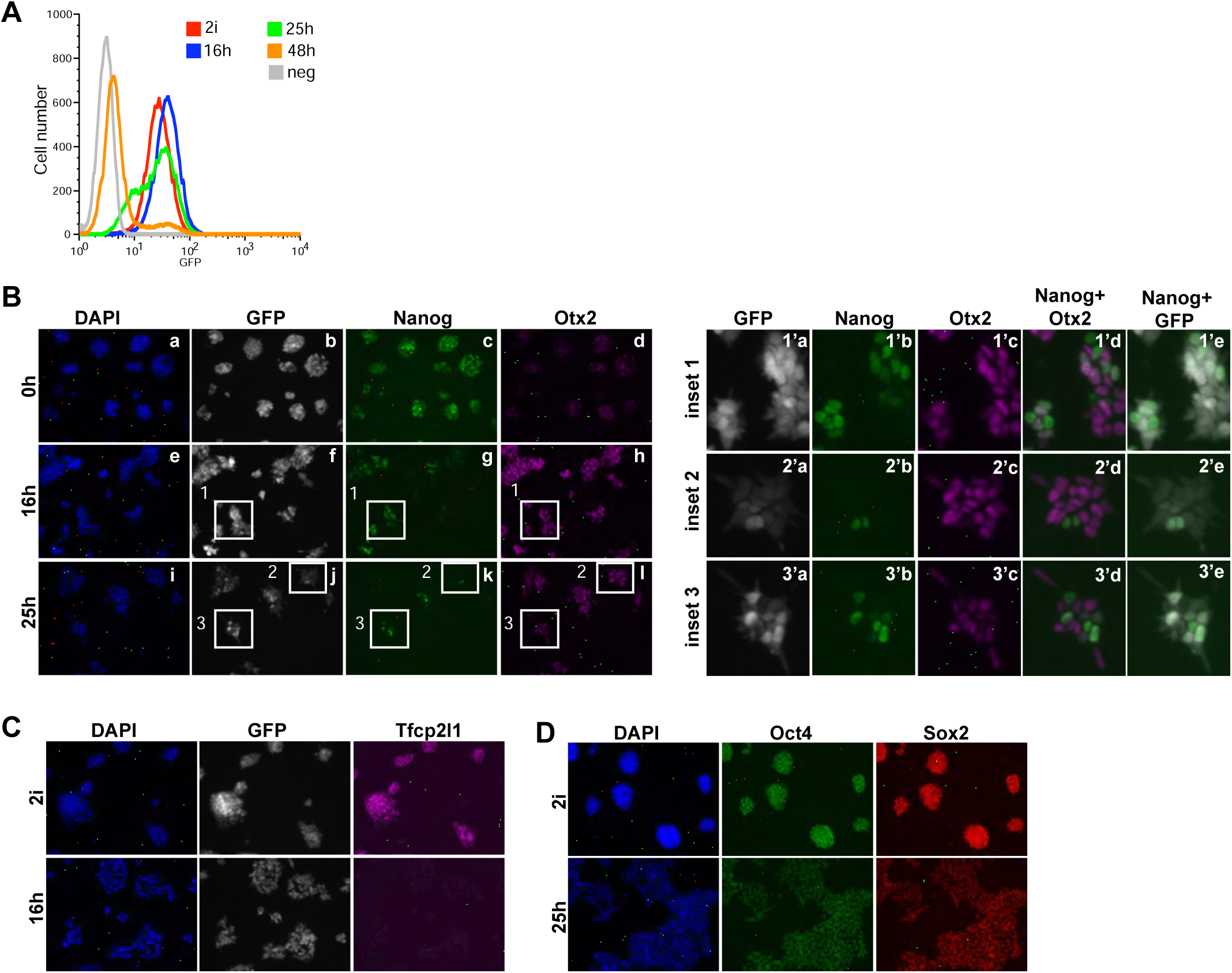
Expression of transcription factors during transition of ES cells. **(A)** Flow cytometry profile for GFP of RGd2 ESCs at indicated time points post-2i withdrawal (from 10000 cells). Wild-type ES cells were used as negative control (neg). IF staining of RGd2 ES cells for **(B)** GFP (gray), Nanog (green), Otx2 (magenta) (a-l). Enlarged versions of the numbered insets are shown in panels a-l. **(C)** GFP (gray) and Tfcp2l1 (magenta) **(D)** Oct4 (green) and Sox2 (red)

We compared the expression of the RGd2 reporter with Nanog and Otx2, a TF that is upregulated in the post-implantation epiblast (Acampora et al., 2013). In 2i Nanog was expressed relatively homogeneously (Fig 3B), while Otx2 was detectable only in a minority of cells (Fig 3C). By 16 h post 2i removal GFP remained ubiquitous, whereas in most of the cells Nanog was downregulated and Otx2 was upregulated (Fig 3B). A few cells co-expressed both TFs, while others showed an exclusive pattern. By 25 h GFP intensity became heterogeneous, consistent with the flow cytometry profile. Occasional Nanog-high cells detected at this timepoint were mostly in the GFP-positive fraction and exhibited low levels of Otx2 (Fig 3B). Conversely, the cells that were strongly positive for Otx2 were mostly negative for Nanog and GFP. A second naive pluripotency factor, Tfcp2l1 (Martello et al., 2013; Ye et al., 2013), was undetectable in the majority of the cells by 16 h (Fig 3C), concomitant with a rapid decrease in mRNA level (Fig 2C). In contrast, Oct4 and Sox2 proteins exhibited homogeneous expression throughout the first 25 h of differentiation (Fig 3D).

These results reveal that TFs associated with pluripotency display individual expression dynamics as ES cells transition from the ground state. RGd2 downregulation appears to track cumulative loss of naive TFs against a background of persistent Oct4 and Sox2 expression.

### Exit from the naive state occurs asynchronously

To determine the time of exit from the naive state, whole populations or subpopulations sorted on the basis of RGd2 expression at selected time points after 2i withdrawal were re-plated at single cell density in serum/L and 2i/L. Resulting colonies were stained for alkaline phosphatase (AP) activity to determine differentiation status (Fig 4A).

**Figure 4.**
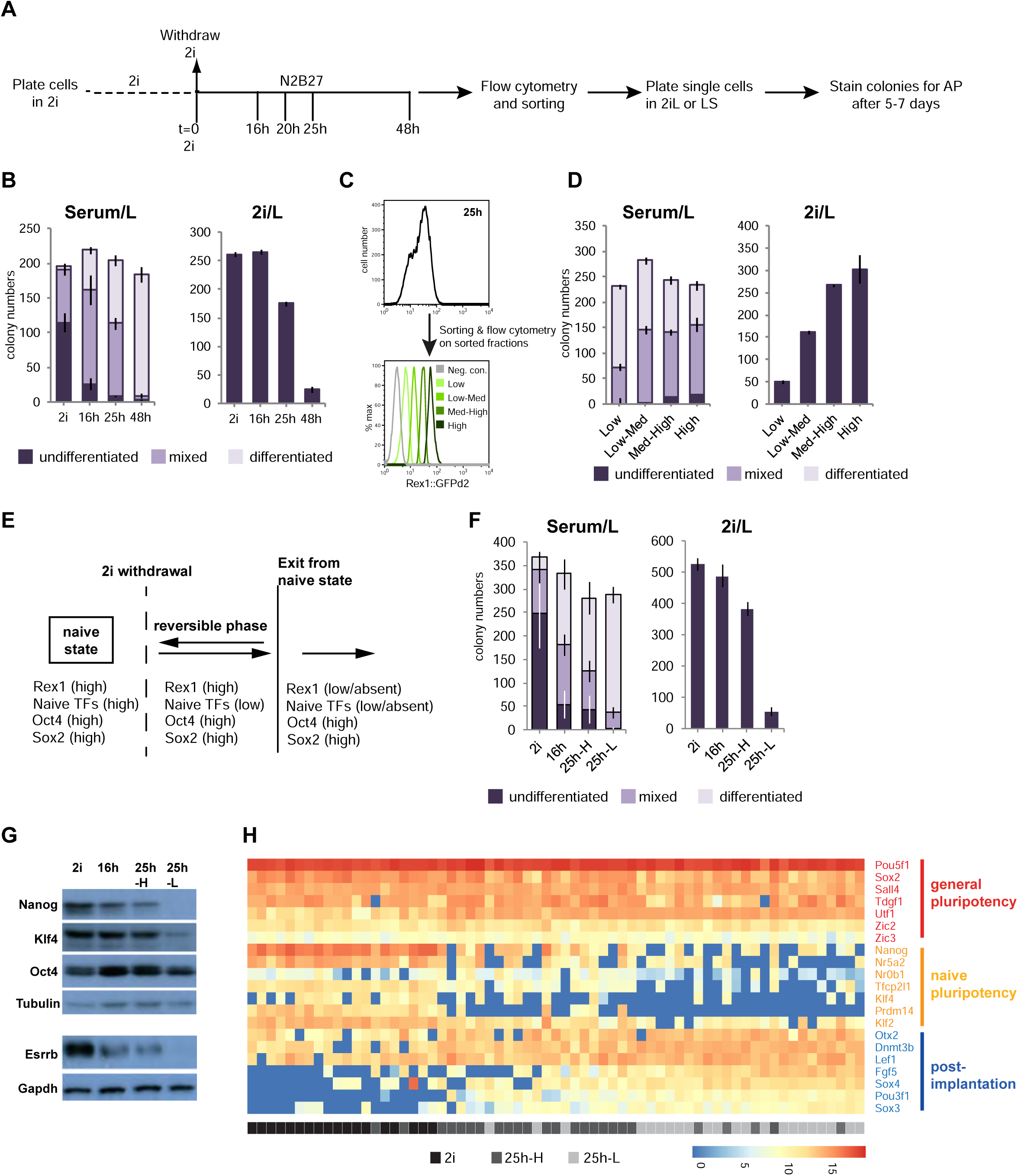
Downregulation of Rex1 tracks exit from the naive state. **(A)** Experimental protocol for cell sorting at indicated time points post-2i withdrawal and clonogenecity assays.**(B)** Clonogenecity of cells from 2i and differentiating subpopulations sorted at indicated time points, plated in serum/LIF (Serum/L) or 2i supplemented with LIF (2i/L). 500 cells per well were plated in duplicate wells of a 6- well plate. Colonies were classified according to alkaline phosphatase (AP) staining performed at day 5 (Serum/L) or day 7 (2i/L) **(C)** Sorting of 25h-cultures into 4 subpopulations based on GFP levels by flow cytometry. Lower plot shows the GFP profiles of post-sort subpopulations **(D)** Clonogenecity of 4 subpopulations shown in **C. (E)** Diagram summarizing phases of transition from the naive state **(F)** Clonogenecity of the indicated subpopulations averaged from 3 independent experiments with 2 technical replicates (related to Fig S2C) **(G)** Immunoblot for indicated proteins on total cell lysates from sorted subpopulations. β-tubulin and Gapdh were used as loading controls. **(H)** Expression levels of selected general (red) and naive (orange) pluripotency and early post-implantation epiblast (blue) markers in single cells from the indicated subpopulations measured by Fluidigm system. Single cells from 2i, 25h-H and 25h-L populations are indicated by black, dark gray and light gray boxes, respectively.

Serum/L permits proliferation of both naive ES cells and differentiating progeny (Marks et al., 2012). Thus, colonies in serum/L reflect plating efficiency and differentiation propensity. Ground state ES cells and cells from inhibitor-withdrawn cultures generated similar numbers of colonies, indicating equivalent plating capacity (Fig 4B). However, the proportions of colony types varied. Even from 2i cells, only around 60% of colonies were wholly undifferentiated, while many were mixed and a few completely differentiated. This heterogeneity is typical of ES cells cultured in serum/L (Wray et al., 2010). The degree of differentiation increased with duration of 2i withdrawal; only 10% of colonies formed after 16 h were undifferentiated while over 20% were wholly differentiated. From 48h-cultures 95% of colonies were differentiated (Wray et al., 2010). Thus ES cells become increasingly predisposed to differentiation in serum as the 2i withdrawal period is prolonged.

In 2i/L self-renewal is optimal, but differentiating cells are eliminated and only naive cells form colonies. Strikingly, the clonogenic efficiency of 16h-cells in 2i/L was equivalent to that of ground state ES cells, even though 16h-cells generated fewer undifferentiated colonies in serum/L (Fig 4B). Thus, the increased tendency for differentiation is not matched by loss of self-renewal potential. However, after 25 h of 2i withdrawal, clonogenecity in 2i/L was significantly reduced, and by 48 h dropped to 10% of naive ES cells. Therefore up to 16 h after 2i withdrawal, self-renewal can be re-asserted, despite the reduction in expression of some naive TFs, induction of postimplantation epiblast markers, and increased tendency to differentiate in serum/L. Between 16 h and 25 h self-renewal capacity is partially lost whereas by 48h exit from the naive state is almost complete across the culture. Thus exit from the naïve state occurs gradually in the ES cell population over an extended period (≤48 h).

### Downregulation of Rex1 tracks exit from the naive state

To determine whether the gradual loss of ES cell identity at the population level is recapitulated at the single cell level we exploited flow cytometry to fractionate cells based on RGd2 expression. We sorted 4 subpopulations 25 h post 2i withdrawal, then replated (Fig 4C, D). In serum/L, colony numbers were relatively constant, although the proportion of undifferentiated colonies declined with decreasing GFP. In 2i/L marked differences in total colony numbers were evident (Fig 4D). The GFP-high fraction exhibited equivalent clonogenicity to 2i cells (Fig 4B, D), indicating complete retention of naive status. However, subpopulations with lower GFP levels produced progressively fewer colonies. The number of colonies formed from GFP-low fraction was only 15% of those formed from 2i or GFP-high cells. Thus the vast majority of this subpopulation has departed the ES cell state (Fig 4D). These data demonstrate that by 25 h the population has become functionally heterogeneous and that cells transition asynchronously.

To examine further whether exit from the naive state and downregulation of Rex1 coincide, we sorted cultures 20 h post 2i withdrawal, when the first GFP-low cells appear, into GFP-high (20h-High) and GFP-low (20h-Low) subpopulations using the same gates used for 25h-cultures. (Fig S2A). Clonogenic efficiency of 20h-High cells in 2i/L was equivalent to ground state ES cells, but was reduced 4-fold for 20h-Low cells (Figs 4B, S2A). Thus the earliest cells that we could obtain after Rex1 downregulation have mostly exited the ES cell state. These data suggest that in individual cells the transition may be relatively acute and occurs at, or slightly after, loss of Rex1 expression.

We examined whether cell cycle asynchrony determines the kinetics of Rex1 downregulation. We stained ES cells with DNA-binding dye Hoechst and isolated subpopulations in G1, S and G2/M by flow cytometry (Fig S2B). We plated these cells, along with stained but unsorted controls, directly in N2B27 at 3x10^4^ cells/cm^2^, which approximates the density at the time of 2i withdrawal in our standard protocol. All populations displayed a similar heterogeneous GFP distribution 25 h after plating, although the G1 starting subpopulation showed a marginally narrower range and a slightly lower mean intensity (Fig S2C). We conclude that the kinetics of Rex1 downregulation is largely unaffected by the initial cell cycle phase.

For subsequent analyses we selected and defined cell populations as follows: 2i-cells represent the ground state; 16h and 25h-H cells are in a reversible phase preceding the loss of ES cell character; and 25h-L cells are the primary products of exit from naïve pluripotency (Fig 4E). Flow cytometry and colony assays confirmed the reproducibility of this system (Figs 4F, S2D, S2E). Immunoblotting showed progressive downregulation of Nanog, Esrrb and Klf4 proteins with time and decreasing GFP, while Oct4 was constant (Fig 4G). The difference between 25h-H and 25h-L cells is of particular note; Nanog and Esrrb proteins are almost undetectable in 25h-L cells and Klf4 is greatly diminished. These three factors are pivotal members of the ES cell gene regulatory circuitry (Dunn et al., 2014) and their elimination together with the absence of Tfcp2l1 (Fig 3C) is expected to be sufficient for loss of self-renewal in 25h-L cells.

To assess further the variation between 25h-cells we performed single cell RT-qPCR for selected genes on 2i, 25h-H and 25h-L populations (Fig 4H). This analysis confirmed that general pluripotency factors remained constant or showed modest changes, whereas naive TFs and post-implantation markers showed variable expression. Heterogeneity was observed for Klf4, Nr0b1, Otx2, Lef1 and Dnmt3b in 2i. Notably, however, all 2i cells were devoid of Fgf5, Oct6 (Pou3f1), Sox3 and Sox4 transcripts, genes that are induced in the post-implantation epiblast (Boroviak et al., 2015). The 25h-H cells showed variable upregulation of these 4 markers and appreciable downregulation of at most 3 naive TFs. In contrast, in 25h-L cells all the post-implantation epiblast markers were induced and at least 4 of the naive TFs were markedly downregulated. These results suggest that decay of ES cell identity during the transition might be defined by the cumulative loss of naive TFs and concomitant gain of factors associated with the post-implantation epiblast. Interestingly, in the reversible 25h-H population, these factors are expressed in various combinations without an evident hierarchy.

### Transcriptional changes during transition from naive pluripotency

To examine global expression dynamics we carried out microarray profiling on 2i, 16h, 25h-H and 25h-L cells, using 3 biological replicates. To directly quantify and confirm gene expression levels, we also performed RNA-seq on independently derived RGd2 ES cell lines. We found a total of 8810 genes in the microarray that were differentially expressed between at least two subpopulations (Table S3). Consistent with the activation of MEK/ERK and GSK3 upon 2i withdrawal, we observed changes in the expression levels of transcriptional targets downstream of these kinases and pathway components. Activation of MEK/ERK is reflected in the upregulation of immediate ERK response genes, such as *Egr1, Fos, myc, c-Jun* (Murphy et al., 2004) and of negative-feedback regulators *Spry2* and ERK phosphatases *Dusp 4* and *6* (Figs 5A). β-catenin functions in ES cell self-renewal via abrogation of Tcf3 *(Tcf7L1)* mediated repression of naïve pluripotency factors (Martello et al., 2012; Pereira et al., 2006; Wray et al., 2011), and in lineage specification through activation of canonical Tcf/Lef1 target genes (ten Berge et al., 2008; Turner et al., 2014; Wu et al., 2012). mRNAs for canonical Wnt targets, T, *axin2, cdx1* and *cdx2* are detected at low levels in 2i, consistent with inhibition of Gsk3 and nuclear accumulation of β-catenin (Marucci et al., 2014) (Fig 5A). Expression is markedly reduced upon 2i withdrawal, indicating reduction of β-catenin dependent transcription during the transition. mRNA levels of transcriptional effectors β-catenin *(Ctnnb1),* Tcf1 *(Tcf7),* Tcf3 *(Tcf7l1)* and Tcf4 *(Tcf7l2)* show modest changes, while Lef1 is significantly upregulated. These observations indicate that loss of ES cell identity is accompanied by increased potential for Lef1-mediated transcriptional regulation.

**Fig 5.**
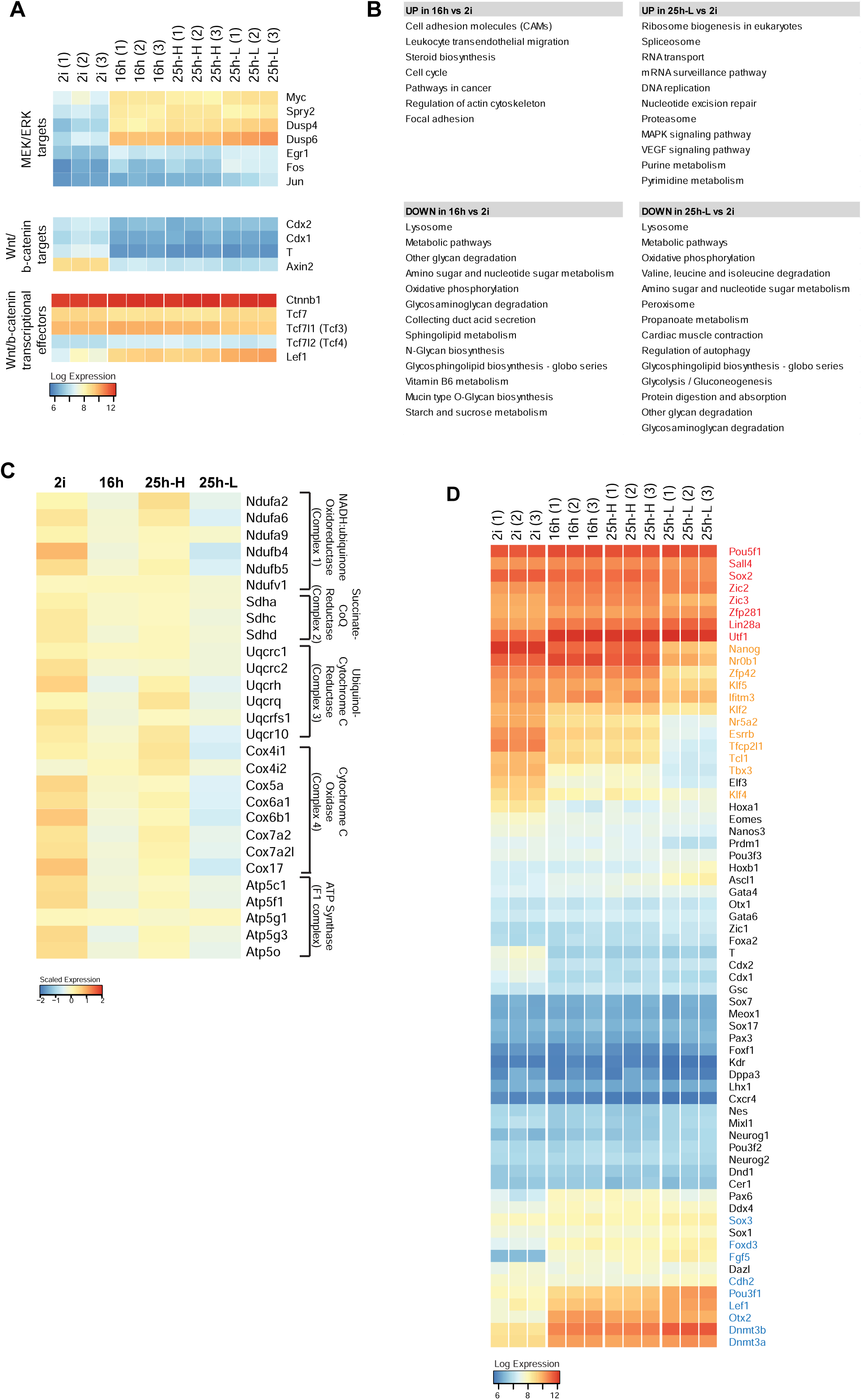
Transcriptional changes in ES cells during progression from naive pluripotency. **(A)** Expression of MEK/ERK and Wnt/b-catenin transcriptional effectors and targets from 3 independent replicates measured by microarray profiling **(B)** Enriched KEGG pathway categories in the differentially expressed gene sets between indicated samples ranked according to p-value (p<0.05) **(C)** Expression levels of differentially expressed mitochondrial ETC complex subunits based on RPKM values measured by RNA-seq **(D)** Expression of general (red) and naive (orange) pluripotency, early post-implantation (blue) and lineage-priming factors (black) measured by microarray.

KEGG pathway enrichment analysis showed that genes with functions in lysosome, oxidative phosphorylation (OxPhos), glycolysis, glycosylation and glycan degradation are the most downregulated in 16h- and 25h-low cells compared to naive ES cells (Fig 5B). Most highly upregulated genes were those associated with cell cycle, cytoskeleton, steroid synthesis and cell adhesion in 16h-cells; and ribosome biogenesis, RNA processing, DNA replication, nucleotide metabolism, proteosome, VEGF and MAPK signalling in 25h-L cells. Further examination of the genes involved in oxidative phoshorylation showed that mRNAs coding for subunits of enzyme complexes that mediate mitochondrial respiration were downregulated in 16h and 25h- H cells, and were further reduced in 25h-L cells (Fig 5C). The changes encompassed all five mitochondrial complexes that mediate mitochondrial electron transport and ATP synthesis. These results suggest that as ES cells leave the naive state, the mitochondrial capacity for oxidative phosphorylation might be diminished. A switch from oxidative to glycolytic energy metabolism during transition from naive to primed pluripotent state has been observed in mouse and human ES cells, and also been proposed to occur during progression of earIy ICM to gastrula stage epiblast (E6.5) (Sperber et al., 2015; Takashima et al., 2014; Zhou et al., 2012). Our analyses indicate that this metabolic transition is initiated prior to lineage priming and might coincide with transition from naive pluripotency.

To benchmark developmental progression, we curated panels of markers for the following categories: general pluripotency; naive pluripotency; post-implantation epiblast; lineage specification (Figs 5D, S3A). The majority of functionally validated naive TFs were reduced significantly in reversible cells, and even further in 25h-L cells, in accordance with decay of ES cell identity. General pluripotency markers either remained relatively constant (Oct4, Sall4, Zic2/3, Zfp281) or exhibited moderate upregulation (Utf1, Lin28b). An exception is Sox2, which was mildly downregulated in 25h-L cells as also seen in the E5.5 epiblast (Boroviak et al., 2015). To assess concordance with protein levels we performed mass spectrometric analysis via stable isotope labelling of amino acids in culture (SILAC). These data confirmed that relative nuclear protein levels of TFs associated with naive and general pluripotency generally correlated with respective transcripts across the different samples (Fig S3B).

Transcripts that are upregulated in the post-impantation epiblast such as Fgf5, Cdh2, Lef1, Otx2, Sox3, Oct6, Foxd3, Dnmt3a and Dnmt3b (Boroviak et al., 2015) were detectable at low levels in 2i cells, but were markedly induced upon 2i withdrawal and highest in 25h-L cells (Figs 5D, S3A). To evaluate whether lineage-restricted factors are expressed, we examined a panel of genes associated with the germ line, neuroectoderm, endoderm or mesoderm. These genes were non-expressed or near background levels in 2i-cells and did not show significant expression levels even in 25h-L cells (Fig 5D, S3C). These results establish that ES cell exit from naive pluripotency is temporally segregated from upregulation of lineage specification factors.

### Comparison of ES cell progression with EpiLC, EpiSC and in vivo epiblast

We compared the RNA-seq data from our ES cell populations (Table S4) to data from embryo samples acquired by a small sample RNA-seq protocol (Boroviak et al., 2015). We also included in this meta-analysis data from post-implantation epiblast-like cells (EpiLC), an intermediate population generated during *in vitro* germ cell differentiation by plating ES cells from 2i/L into Fgf2, Activin and 1% KSR for 48 h (Buecker et al., 2014; Hayashi et al., 2011). Principle component analysis was based on 1857 dynamically expressed genes between E2.5 and E5.5 (Boroviak et al., 2015) Separation between *in vitro* and in vivo samples may largely be attributed to different RNA-seq methods because differently processed ES cell samples cluster apart according to protocol (Fig 6A). The embryo samples resolve according to developmental stage. ES cell-derived samples are less separated but align with the developmental trajectory from E4.5 epiblast towards E5.5 post-implantation epiblast. The reversible 16h and 25h-H populations cluster closely together while the 25h-L population is more advanced and EpiLC lie between. EpiLCs are formed in the presence of exogenous inducers and differential gene expression analysis revealed a number of genes that distinguish them from 25h-L cells (Table S5). We also generated EpiLC from RGd2 ES cells and measured reporter expression by flow cytometry. We found that a subpopulation of EpiLCs (23%) express Rex1 at naive ES cell levels (Figs S4A, B), indicating that EpiLC populations may contain residual undifferentiated ES cells.

**Fig 6.**
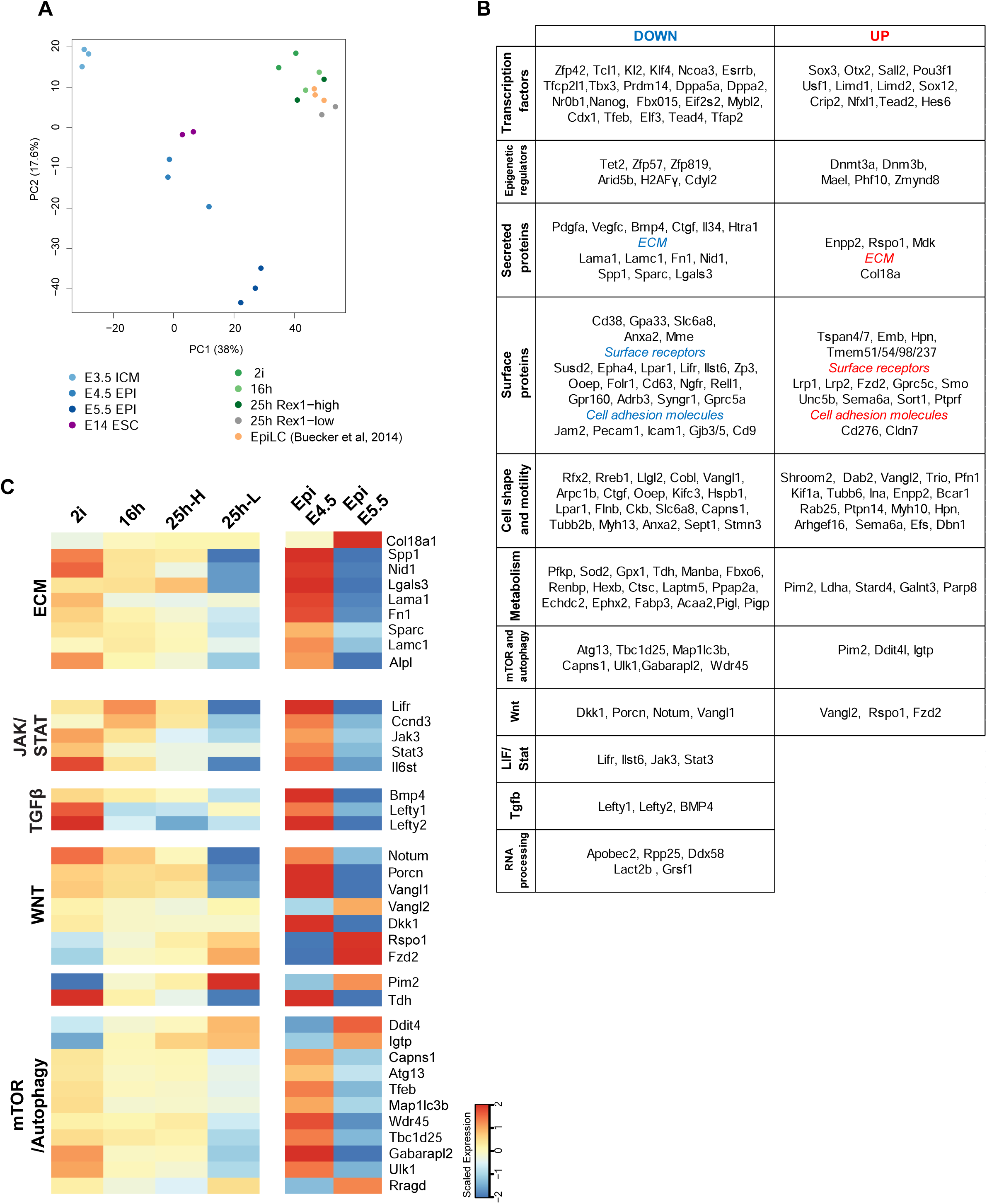
Comparison of transcriptional changes during pluripotency progression in ES cells and in the embryo. **(A)** Principal component analysis of RNA-seq datasets based on dynamically expressed genes between E2.5 and E5.5 **(B)** Functional grouping of genes that show similar regulation in ES cells and in the embryo **(C)** Expression of genes from selected pathways

We additionally undertook a comparison with published data on EpiSC that are related to E6.5 epiblast (Kojima et al., 2014). A heatmap of marker expression (Fig S4C,D), shows that 25h-L cells are related to E5.5 epiblast and are distinct from EpiSC. These data confirm that ES cells do not transition directly into EpiSC (Hayashi et al., 2011).

To examine the embryo correspondence further we isolated expression changes that occur between E4.5 and E5.5 as the epiblast cells exit from naive pluripotency. We asked to what extent these changes are reflected in the *in vitro* transition. Out of 608 differentially expressed genes in the epiblast and robustly detected in at least one of the samples (fragments per kilobase of exon per million fragments mapped [FPKM] ≥10), 190 do not change expression appreciably between 2i ES cells and 25h-L cells, while 52 showed differential expression in the opposite direction to the embryo (Table S6). This latter subset included ERK target Egr1 and factors that regulate cell proliferation (Atf3, Tef, Trp53, Tada3, Klf6, Ccng1), apoptosis (Apaf1, Bid), cell adhesion and morphology (Krt18, Cdh2, Fez1, Lamb1, Tns3, Amotl1) as well as signalling pathway components, such as, Notch receptor Notch3 and Notch nuclear effector Rbpj and Nodal co-receptor Tdgf1. Contrasting regulation of these genes might reflect differences in the duration and intensity of signalling responses downstream of MEK/Erk and GSK3 *in vitro,* or the absence of paracrine signalling cues in the minimal culture system. However, the majority (366 of 608) of genes that are differentially expressed between E4.5 and E5.5 exhibited differential expression during ES cell transition, with the direction of change conserved (Fig 6B, C, Table S6). Several functional groups could be identified within the common up- and down- regulated gene sets (Fig 6B). Besides transcription factors and epigenetic regulators with established functions in pre- and early post-implantation development, the sets included genes associated with extracellular matrix (ECM), cell adhesion, motility, shape, metabolism and autophagy. Notably, mRNAs for structural ECM components Fibronectin (Fn1), Laminin isoforms, Lamc1 and Lama1, Laminin-linker protein Nid1, Spp1 (Osteopontin), Sparc, Lgals3 were downregulated (Fig 6C). Interestingly, non-tissue specific alkaline phosphatase (Alpl) that is used as a surrogate marker for ES cells was also among the downregulated genes. Alpl modifies the ECM by dephosphorylating ECM molecules such as Osteopontin (Spp1) (Narisawa, 2015). Conversely, Col18a was upregulated indicating major reconfiguration of ECM during exit from naive pluripotency.

Another common feature of *in vitro* and *in vivo* transitions is downregulation of LIF receptor components, Lifr and Ilst6, signal transducers Jak3 and Stat3 and the transcriptional target Ccnd3, indicating that responsiveness to LIF is dimininished as cells transition from naive pluripotency (Fig 6C). Tgfβ ligand BMP4 and Nodal inhibitors, Lefty1 and Lefty2 were also downregulated, suggesting modulation of the Tgf-β signalling output during the transition. Additionally, potential modulation of Wnt signalling was reflected by changing levels of its positive and negative regulators.

We also noticed changes in the mRNA levels of enzymes that regulate and respond to cellular energy metabolism. Threonine dehydrogenase (Tdh) was dramatically downregulated. Tdh is vital for ES cell survival for its function in conversion of threonine into acetyl co-A and glycine, providing precursors for mitochondrial TCA cycle and purine synthesis (Wang et al., 2009). In contrast *Pim2* is one of the most highly upregulated genes in both settings (Fig 6C). Pim2 can promote glycolysis and mTOR signalling by directly phosphorylating pyruvate kinase M2 (PKM2) (Yu et al., 2013) and TSC2, respectively, relieving repression of mTOR-C1 (Lu et al., 2013; Zhang et al., 2015). Other genes that regulate or respond to mTOR signalling also showed increased expression. mTOR pathway inhibitor Ddit4 (Redd1), and mTOR activator Rragd were upregulated, suggesting modulation of mTOR activity. Activated mTOR suppresses autophagy by phosphorylating and inhibiting Ulk1 and Atg13, two molecules that are required for autophagy initiation, and through phosphorylation-dependent cytoplasmic sequestration of Tfeb, a key TF that orchestrates expression of genes involved in lysosome function and autophagy (Kim and Guan, 2015; Napolitano and Ballabio, 2016). These three mTOR targets, Ulk1, Atg13 and Tfeb, along with several autophagosome- associated factors were downregulated in both settings (Fig 6C), suggesting transcriptional and post-transcriptional suppression of autophagy as the cells transition from naive pluripotency.

Collectively these transcriptome analyses support the idea that loss of Rex1 expression in a defined ES cell culture system reflects features of the developmental transition from the pre- to post-implantation pluripotency.

### Acquisition of DNA methylation during transition from naïve pluripotency

Genome-wide DNA methylation increases substantially between E4.5 and E5.5 in utero (Auclair et al., 2014). Expression of *de novo* DNA methyltransferases Dnmt3a and Dnmt3b is markedly upregulated, whereas Prdm14, which represses Dnmt3a/b and promotes Tet activity on target genes (Ficz et al., 2013; Okashita et al., 2014; Yamaji et al., 2013), is downregulated both in ES cells and in the epiblast during transition (Figs 7A, 5D, S3A). We therefore performed whole genome bisulfite sequencing on naïve ES cells and GFP-sorted transitional populations. We detected an increase in total CG methylation across the genome upon 2i withdrawal (Fig 7B). Notably, average genome methylation tended to increase in small increments between 2i, 16h and 25h-H, but a pronounced and statistically significant increase was observed for 25h-L. The level of increase was similar across gene bodies, exons, introns, intergenic regions, satellites and retrotransposon sequences (LINEs, SINEs, LTRs, IAPs), while methylation of CpG islands (CGIs) was not generally altered (Fig S5A, B). Promoters that contain CGIs were hypomethylated in 2i and remained refractory to DNA methylation upon 2i withdrawal, while promoters without CGIs (non-CGI-promoters) exhibited increased methylation similar to the genome average (Fig 7C).

**Fig 7.**
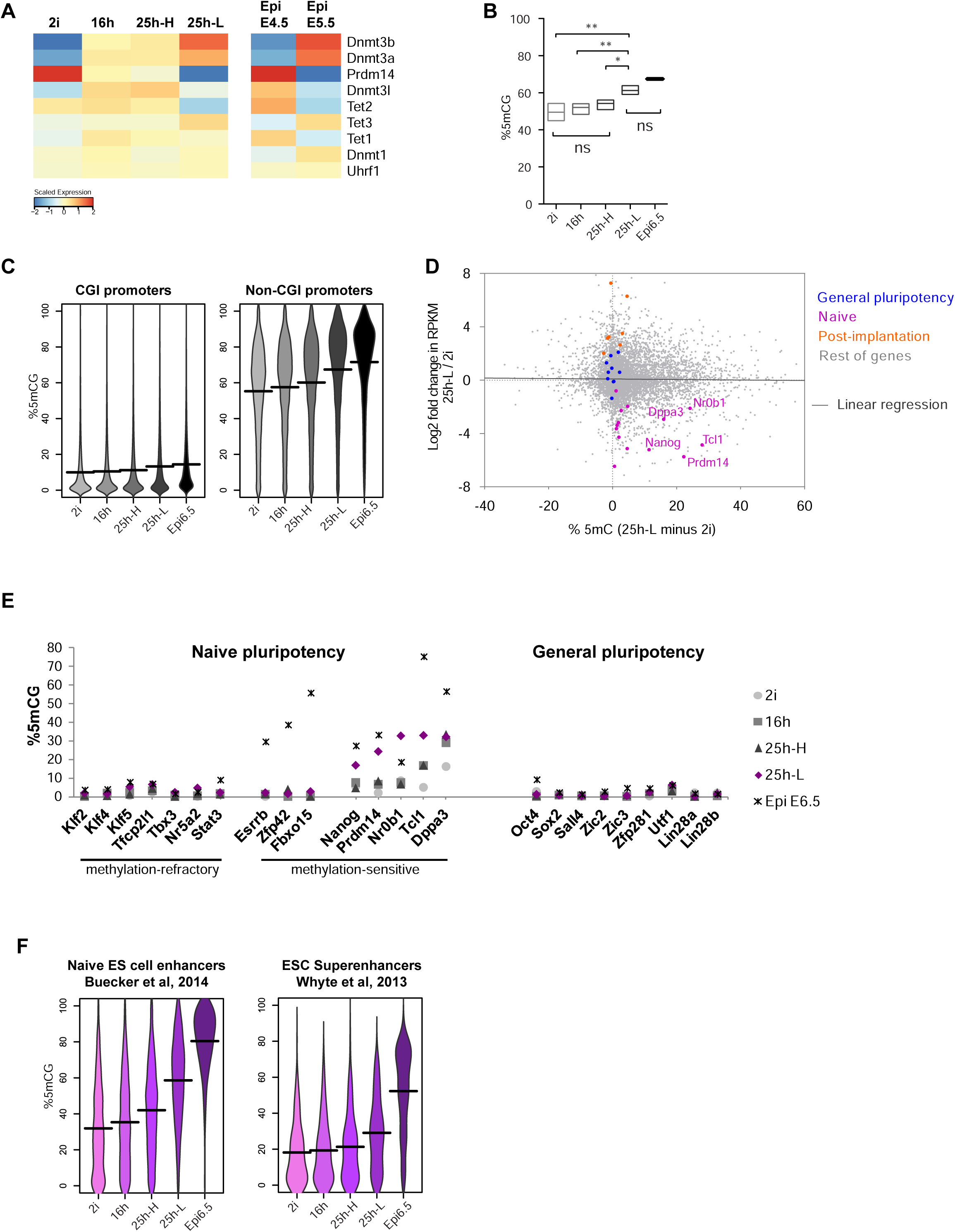
Acquisition of DNA methylation during transition from naïve pluripotency. **(A)** Expression of factors that modulate DNA methylation **(B)** Global genomic methylation in CG context (mCG) **(C)** Percentage of mCG in the promoters of expressed genes **(D)** Comparison of changes in the transcript levels with changes in promoter methylation between 2i and 25h-L cells. Coloured dots represent individual genes belonging to 3 groups: general pluripotency (blue), naive pluripotency (pink), post-implantation (orange), as in heatmap in Fig 5D. Black line shows linear regression. Levels of mCG in **(D)** the promoters of naive and general pluripotency genes **(F)** enhancers.

Whole genome methylation data are not available for E5.5 epiblast. We therefore compared our profiles with published data on E6.5 post-implantation epiblast (Seisenberger et al., 2012). This analysis showed that the CG methylation level of 25h- L cells is intermediate between naïve ES cells and E6.5 post-implantation epiblast across all genomic regions (Figs 7, S5). This is in line with reduced representation bisulfite sequencing data on embryos showing a marked increase in methylation between E4.5 and E5.5 with a further increase at E6.5 (Auclair et al., 2014).

To determine if promoter methylation correlates with transcript levels, we examined the promoters of expressed genes (RPKM≥10 in at least one population). We plotted the difference in percentage CG methylation of the promoters against the fold change in transcript levels between 2i and 25h-L cells (Fig 7D). Although there was no overall anti-correlation (R^2^ = 0,0005744), we found that gains in DNA methylation in a subset of naïve pluripotency gene promoters were associated with decreased transcript levels, including for *Nanog, Prdm14, Nr0b1, Tcl1, Dppa3.* Promoters of naive and general pluripotency genes were almost completely unmethylated 2i, with the exception of *Dppa3* which displayed low level methylation (Fig 7E). Upon 2i withdrawal, the aforementioned *Nanog* subgroup acquired methylation, most notably in 25h-L cells, albeit less than in the E6.5 epiblast. However, promoters of general pluripotency genes and the other naive pluripotency genes remained unmethylated. These latter promoters are unmethylated also in the E6.5 epiblast, with the exception of *Esrrb, Zfp42* and *Fbxo15* (Fig 7E). Thus gain of methylation is not entirely random and pluripotency-associated genes acquire methylation with different kinetics.

To investigate further how DNA methylation might relate to gene expression we examined enhancers. We identified potential “naïve enhancers” from published ChIP-seq datasets (Buecker et al., 2014) as regions displaying the general enhancer mark H3K4me1, together with the active enhancer marks H3K27Ac and p300 in naïve ES cells (Blow et al., 2010; Heintzman et al., 2007; Rada-Iglesias et al., 2011; Visel et al., 2009; Visel et al., 2013). In 2i these enhancers were under-methylated compared to the genome average, but they gained methylation progressively upon 2i withdrawal (Fig 7F). We also examined the methylation state of ES cell “super enhancers” (SEs) defined in serum/L (Hnisz et al., 2013; Whyte et al., 2013). The SEs exhibited low and relatively constant methylation levels in 2i, 16h and 25h-H cells, with a small increase in 25h-L cells that remained below the genome average (Fig 7F). These observations indicate that accompanying exit from naïve pluripotency ES cell enhancers associated with naïve pluripotency are methylated, indicative of decommissioning, while SEs linked to general pluripotency-associated transcription are protected from methylation.

## Discussion

Our results demonstrate that the RGd2 reporter enables near-real time detection of exit from naive pluripotency and isolation of the first cells to alter functional state. Loss of Rex1 expression marks a change in pluripotent identity that precedes a decline in Oct4 level or acquisition of lineage-specific gene expression. The defined *in vitro* monolayer differentiation system lacks uterine or extraembryonic cues that are considered to guide epiblast development in the embryo. In these minimal conditions autocrine signals are sufficient to drive transcriptome and methylome changes that are generally convergent with peri-implantation epiblast. These findings indicate that the gene regulatory circuitry of ES cells has an innate capacity to orchestrate a profound developmental transition.

At the onset of this transition, the molecular network that sustains naïve pluripotency is dismantled (Buecker et al., 2014; Kalkan and Smith, 2014; Leeb et al., 2014). Apparently co-incident with acute downregulation of the critical naive TFs, postimplantation epiblast markers are up-regulated. Increased differentiation when transferred to serum suggests enhanced sensitivity to inductive cues even before cells have fully extinguished ES cell identity. However, for as long as Rex1 is expressed, cells retain in full the ability to regain the ES cell ground state. Such reactivation of self-renewal, despite marked reduction in the levels of functionally important naïve TFs, is consistent with the postulate that the ES cell state is founded on a highly flexible transcription factor circuitry (Dunn et al., 2014; Martello and Smith, 2014; Niwa, 2014; Young, 2011). We surmise that loss of Rex1 reflects a cumulative reduction of the suite of factors below a critical threshold. From this point the network cannot be reactivated and is subsumed by the emerging new network. A gradual loss of selfrenewal apparent from whole population analyses reflects asynchronous single cell dynamics and at the level of individual cells the transition may be quite acute.

Rex1 transcription is considered to be directly regulated by several naïve TFs (Chen et al., 2008; Kim et al., 2008) which can explain how the RGd2 reporter serves as a sensor of the overall activity of the naïve transcription factor circuitry. Nevertheless, 10-15% of the Rex1-low cells at 25h can be restored to ground state self-renewal. This may be explained in part by incomplete efficiency of flow sorting, but also suggests that Rex1 downregulation might be separated from exit from the naïve state in a minority of cells. Indeed the higher incidence of reversion for Rex1-Low cells at 20h is consistent with silencing of Rex1 slightly preceding loss of ES cell identity. Alternatively the connection between down-regulation of naïve factors and loss of Rex1-GFP may not be purely deterministic. Rare Nanog-positive/GFP-negative cells detected at 25h could be consistent with the latter explanation. Fluctuating Rex1 reporter expression has been reported in serum (Toyooka et al., 2008), where ES cells are continuously exposed to conflicting pro- and anti-differentiation stimuli that may perturb developmental progression. Even in those conditions, however, it is apparent that with more stringent categorisation of Rex1-negative cells, reversion frequency is diminished (Nakai-Futatsugi and Niwa, 2016). Moreover, Rex1-negative cells in serum culture tend to be eliminated from blastocyst chimaeras (Alexandrova et al., 2016; Toyooka et al., 2008).

Consistent with loss of functional ES cell character, the 25h-L population show significant transcriptome variance from their naïve predecessors. They are clearly distinct from EpiSC, however, and appear to be converging towards E5.5 epiblast. Currently comprehensive data from this stage in utero is lacking and it will be of great interest to examine this relationship further as new data become available, and also to determine to what extent microenvironmental modulations, such as substrate composition, may further increase the veracity of the ES cell model.

The dynamic and global increase in DNA methylation as ES cells leave the naïve state generate a methylome intermediate between naïve and primed pluripotent compartments *in vitro* and *in vivo.* Interestingly, most pluripotency gene promoters were spared from methylation. Among the exceptions that gained methylation, however, are *Nanog* and *Prdm14.* Both factors may repress Dnmt3b directly and have been proposed to recruit Tet enzymes to target loci (Costa et al., 2013; Ficz et al., 2013; Okashita et al., 2014). Moreover, Prdm14 represses expression of Fgf pathway components (Yamaji et al., 2013). Thus, silencing of Nanog and Prdm14 through promoter methylation might render 25h-L cells unable to reverse DNA methylation events or signalling activity. In the context of transcriptional down-regulation of naïve TFs and loss of LIF-responsiveness, these effects, along with methylation of naive enhancers, may present an additional barrier to re-entry to the naïve state.

We have proposed that downregulation of naïve pluripotency factors elicits differentiation competence via an intermediate phase of “formative pluripotency”. This is postulated as a period of competence acquisition for multi-lineage specification (Kalkan and Smith, 2014). *In vivo* the formative phase corresponds to peri- and immediate post-implantation epiblast (E4.75-5.75), before cells exhibit expression of lineage specification factors. Notably during this period epiblast cells acquire competence for germ cell induction (Hayashi et al., 2011; Ohinata et al., 2009). Our results indicate that ES cells that downregulate Rex1 and depart naïve pluripotency show transcriptome and methylome features that may be anticipated for immediate post-implantation epiblast. Thus, these Rex1-low cells represent a snapshot of the nascent formative phase, undergoing rewiring of the gene regulatory network and remodelling of the methylome. The datasets we provide constitute a resource for inspecting an overlooked phase of pluripotency. It will be of future interest to dissect in detail the molecular dynamics and drivers of transition in this defined and simple system and also to determine whether the formative phase may be suspended in culture, as for naïve ES cells and primed EpiSCs.

## Acknowledgements

This research was funded by the Wellcome Trust, the Biotechnology and Biological Sciences Research Council, European Commission Framework 7 project EuroSyStem, and the Louis Jeantet Foundation. The Cambridge Stem Cell Institute receives core funding from the Wellcome Trust and Medical Research Council (MRC). AS is an MRC Professor.

## Materials and Methods

### Mouse colony establishment and immunostaining of embryos

Mice were maintained as described previously (Nichols et al., 2009a). RGd2.c6 ES cells carrying a GFPd2-IRES-Blastcidin expression cassette between the translation start and stop codons of one of the *Zfp42* (Rex1) alleles (Wray et al., 2011) were injected into E3.5 C57Bl/6 blastocysts. Offspring were assessed for chimaerism by coat colour. Three male chimaeras with a high degree of coat colour contribution were bred with wild-type 129 females. The resulting offspring that genotyped positive for the Rex1-GFPd2 allele was back-crossed to wild-type 129 animals once more. Following this, heterozygous offspring were crossed to generate homozygous mice. Homozygous mice were then bred to generate a stock of mice homozygous for RGd2 reporter. Immunostaining was performed as described previously (Nichols et al., 2009b) using antibodies listed in Table S7. Embryos were imaged on a Leica TCS SP5 confocal microscope.

### ES cell lines and culture

ES cell lines carrying the RGd2 reporter were derived from embryos using previously described protocols (Nichols et al., 2009a). For routine maintenance ES cells were plated at 2-3 x 10^4^ cells cm-^2^ in 2i on 0.1% gelatine-coated dishes (Sigma, cat. G1890) and passaged every 3 days following dissociation with Accutase (PAA, Cat. L11-007). 2i consists of N2B27 (NDiff N2B27 base medium, Stem Cell Sciences Ltd, Cat. SCS-SF-NB-02) or homemade N2B27, supplemented with PD0325901 (1 μM) and CHIR99021 (3 μM). LIF prepared in house was added only when indicated.

### Immunoblot and immunofluorescent staining of ES cells

ES cells were lysed in lysis buffer [1xPBS, 1%TritonX-100, 0.1%SDS] with protease and protein inhibitors (Roche) and sonicated briefly in the Bioruptor (Diagenode) to shear the gDNA. Total cell lysates equivalent to 2-4x 10^4^ cells were loaded per lane. Primary antibodies and dilutions used are listed in Table S7. HRP-conjugated secondary antibodies and ECL reagent (GE Healthcare Life Sciences) were used for detection. For immunofluorescent (IF) staining, cells were fixed for 10min with 4%PFA at RT, followed by permeabilization and blocking in PBS containing 0.1% TritonX-100 and 3% donkey serum. Cells were incubated with primary antibodies in blocking solution overnight at 4C. Alexa Fluor-conjugated donkey secondary antibodies (Molecular Probes) were used at 1:1000.

### Monolayer differentiation, cell sorting and clonogenecity assays

For monolayer differentiation in N2B27 cells were plated at 1.5 x 10^4^ cells cm-^2^ in 2i and medium was replaced with N2B27 after 20-24 h. Prior to sorting, cells were pelleted and resuspended in the previous culture medium. For dissociation of 2i-cells 2i inhibitors was added to Accutase. ToPro-3 (Invitrogen) was added at a concentration of 0.05 nM to label membrane compromised cells. Cells were sorted in MoFlo flow sorter (Beckman Coulter). For analysis of 2i -and 16h-cultures, all ToPro-3-negative cells were collected. For sorting Rex1-high and -low cells from 25h-cultures, gates were set to isolate subpopulations with highest and lowest 15% GFP expression. For clonogenecity assays, 500-800 cells were plated on 6-well dishes in L/S or 2i/L, coated with 0.1% gelatine or laminin (Sigma, Cat. L2020), respectively. On day 4 (L/S) or 6 (2i/L), plates were fixed and stained for AP (Sigma, Cat. 86R-1KT). Plates were scanned using a Cell Celector (Aviso) and scored manually. Colony formation efficiency for a given population was determined by dividing the averaged number of colonies formed in 2i/L by that in L/S. For detection of RGd2 reporter expression, flow cytometry analysis was performed using a Dako Cytomation CyAn ADP high-performance cytometer and the results were analyzed with Summit software.

### RNA extraction, cDNA synthesis and qPCR

For all RNA analyses, total RNA was isolated using RNeasy mini kit (Qiagen). cDNA was synthesized using SuperScript III (Invitrogen) and oligo-dT primers. qRT-PCR was performed with TaqMan Gene Expression assays (Thermo Scientific) using probes listed in Table S7.

### Single cell qRT-PCR

Cells were sorted using a G1 enrichment strategy, based on forward scatter (FS) and side scatter (SC) gating. Single cells were sorted into 96-well plates containing CellsDirect One-Step qRT-PCR master mix (Invitrogen, 11753-100) for cDNA synthesis and pre-amplification. Samples were shipped to the Genomics Core Facility at the European Molecular Biology Laboratory in Heidelberg, Germany, where Fluidigm assays were run according to the manufacturer's protocols using EvaGreen detection and primer sets listed in Table S7.

### Microarray, RNA-sequencing, DNA methylome and proteome analyses

Processing of ES cell samples and data analyses are described in the Supplementary file.

